# Understanding the Sport Viewership Experience using Functional Near-Infrared Spectroscopy

**DOI:** 10.1101/2024.08.01.606260

**Authors:** Luke R. Potwarka, Adrian B. Safati, Adam T. Pappas, Girish Ramchandani, Michael L. Naraine, Nur Gurbez, Peter A. Hall

## Abstract

Subjective evaluation of a sport event in real time is normally assessed using self-report measures, but neural indices of evaluative processing may provide new insights. The extent of evaluative processing of a sporting event at the neural level may depend on the degree of emotional investment by the viewer, as well as the key moment of the game play being observed. Those with high ego involvement might show more activation within evaluative processing nodes, and this pattern may be most pronounced during critical moments of game play. In the current study, we examined neural activations within the medial and lateral prefrontal cortex during game play as a function of ego-involvement, using video clips featuring key moments in a European league ice hockey game. A total of 343 participants were pre-screened to identify 20 high and low ego-involved individuals. These subgroups then viewed a game segment containing 12 key play moments, while undergoing neuroimaging using fNIRS. Findings indicated more engagement of the dmPFC throughout all key moments for high ego-involved participants, but particularly during critical game moments. Overall, findings suggest that neural indices of evaluative processing might contribute meaningfully to understanding when emotionally invested individuals are most engaged in an action sequence during a sporting event.

## Introduction

Theories and models of sport consumer behavior often rely on self-reported measures when attempting to understand spectator/viewer experiences (Funk & James, 2001; Madrigal, 2006). Self-report measures are subject to retrospective biases and lack in-situ perspectives; as well, thoughts and feelings operate at a subconscious-level, and this may not manifest in subjective impressions (Oppenheim, 1992). Neuroimaging techniques provide researchers with new ways of understanding brain-related mechanisms underlying spectator experiences in a more direct manner (Potwarka et al., 2022), but they are also challenging to implement in the context of *in vivo* sporting events. Partially as a function of this, sport consumer researchers have rarely explored how subjective impressions of sport viewership experiences might relate to patterns of brain activation, or how what kinds of critical moments within sporting events elicit specific types of brain activity. Functional near-infrared spectroscopy (fNIRS) is a promising solution because of its flexibility and adaptability for in-vivo measurements of brain activity, particularly those that require upright viewing to enhance ecological validity (Ayaz et al., 2022; Ferrari & Quaresima, 2012; Potwarka et al., 2022). Consistent with Thomson et al. (2019), we advance the position that measuring brain activity of consumers in the inherently social context of sport event viewing provides access to psychological processes and neural circuitry that guide engagement.

One self-report construct of interest to sport consumer researchers is that of ego-Involvement (EI), an unobservable state of motivation, arousal or interest toward a sport activity, event, and/or associated product (Kyle et al., 2007; Rothschild, 1984). It is evoked by a particular stimulus or situation and has drive-like properties (Rothschild, 1984). Typically, EI is measured in terms of how central a sport is to one’s life; and the degree to which the sport (or related behavior or experience) is connected to one’s identity (Kyle at al., 2007). EI is an important construct in sport consumer research because it has been linked to loyalty, repeat purchase behaviour and satisfaction (Furley et al., 2013; Havitz & Dimanche, 1999). Moreover, the EI construct underpins several widely used models of sport consumer behavior and major theoretical perspectives within sport (e.g., Funk & James, 2001).

The purpose of the present investigation was threefold. First, we wanted to examine the viability of fNIRS as a method for assessing neural activity in an ecologically valid, *in-vivo* sport viewership experience. Second, we were interested in examining the extent to which different key moments within a sport event were differentially predictive of changes in brain activity within the neocortex. Finally, we endeavored to examine the extent to which baseline identification with the sport (i.e., ego-involvement) moderates the above effects. We specifically hypothesized that, beyond fNIRS being a well-suited method for the sport viewership context, cerebral hemodynamics would reveal increases in activation during pivotal game play moments, that that these effects would be particularly pronounced for those with relatively higher levels of ego-involvement prior to the viewership experience.

*A-priori* regions of interest included the left and right dorsomedial prefrontal cortex (dmPFC) because of their links to evaluative and self-relevance processing (Etkin et al., 2011; Lieberman, Straccia, Meyer, Du & Tan, 2019; Meyer & Lieberman, 2018), which could be disproportionately engaged among ego-involved viewers relative to their more neutral counterparts. In addition, the left and right lateral prefrontal cortex (lPFC) were chosen because of links with attentional engagement (Kam et al., 2018; Kondo et al., 2004), which could arguably be stronger among those with higher ego-involvement. Heighted activity in the dmPFC could reflect viewers’ emotional involvement in the game action, feeling a sense of connection with a team, or experiencing personal relevance in relation to the game’s outcome. Lateral PFC activation in these areas is indicative of working memory and top-down attentional control-filtering out distractions (Harris et al., 2013). These processes are essential for following game progress and trying to make sense of the gameplay as it unfolds.

## Methods

### Participants

Study participants were 20 young adults between the ages of 18 and 40 (*M* = 21.84, *SD*=5.08). A table showing demographic characteristics of the sample and study variable means and variability indices is shown below (Table 1).

**Table 1:**
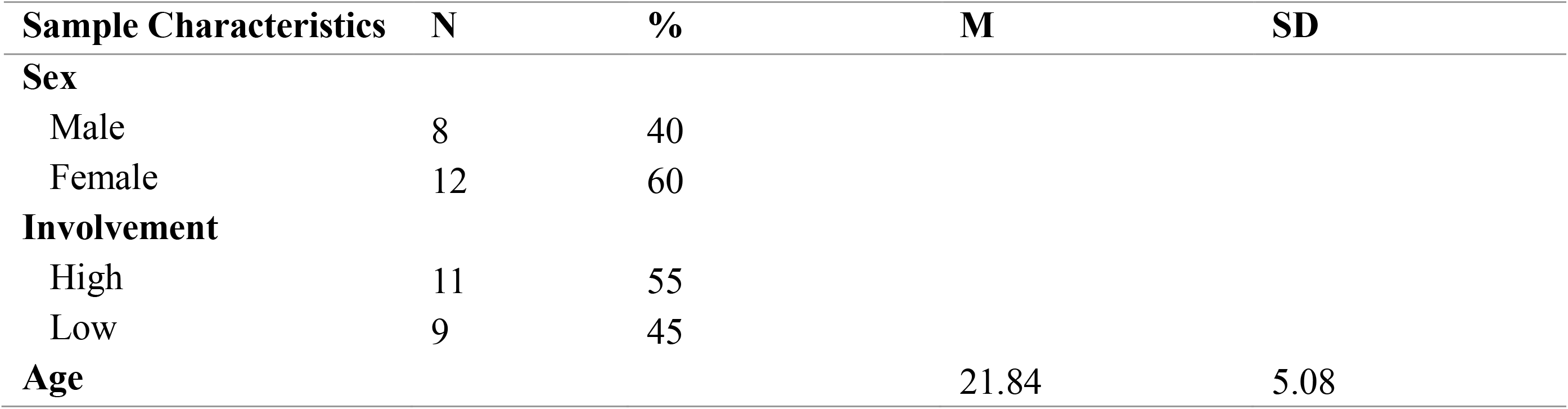
Demographic Characteristics of the Sample.

| Sample Characteristics | N | % | M | SD |
| --- | --- | --- | --- | --- |
| **Sex** |  |  |  |  |
| Male | 8 | 40 |  |  |
| Female | 12 | 60 |  |  |
| **Involvement** |  |  |  |  |
| High | 11 | 55 |  |  |
| Low | 9 | 45 |  |  |
| **Age** |  |  | 21.84 | 5.08 |

### Procedures

In an initial screening phase, an in-person survey was distributed to undergraduate students (*n*=343) attending classes at a large University in Southern Ontario, Canada. The questionnaire included a measure of EI in relation to watching ice-hockey; this measure was used to stratify potential respondents in terms of *a-priori* engagement and interest in the target behavior. In the laboratory phase of the study, 20 individuals were required from the “high involvement” (*n*=11) and “low involvement” (*n*=9) subgroups, and subsequently invited to join a laboratory session.

Upon entry to the lab, participants underwent an informed consent procedure and were seated at a laptop computer. The fNIRS measurement band (containing 16 optodes) was fitted to the forehead, and participants viewed a segment of ice hockey play from the Elite Ice Hockey League (United Kingdom) game between the Nottingham Panthers (wearing black) and the Cardiff Devils (wearing red). Prior to watching the game, participants were instructed to choose a team to support. This league and set of teams were chosen because the participants were unlikely to have had prior experience viewing gameplay involving the targets. The play segment (the first 20-minute period of the ice hockey game) was a video clip lasting approximately 25 minutes, including play breaks. At each of the scoring chances and game stoppages (offensive faceoffs), a slice of video segment was cut and presented as stimuli; there were 12 such segments used as stimuli in the imaging portion of the study described below.

### Measures

#### Cortical hemodynamics

Functional near-infrared spectroscopy (fNIRS) uses near-infrared spectrum light emitters paired with sensitive light detectors (optodes) to measure subtle changes in oxygenated hemoglobin concentration (HbO) within the human neocortex (Pinti et al., 2020). Task-related hemodynamic responses were assessed using an fNIR Devices unit 203c. To measure cortical activation, a fabric band containing the fNIRS optodes was placed over the forehead using anatomical landmarks and fastened with Velcro fasteners. The optodes consisted of 4 light sources and 8 detectors, which combined could produce up to 16 measurement channels, with a sampling rate of 5Hz. Both oxy- and de-oxyhemoglobin were measured. For the purposes of this study, oxyhemoglobin was measured as the primary metric to quantify neural activation. *A-priori* slices of play action (described above; *n*=12 were played on a computer screen in front of each participant while hemodynamic changes in the prefrontal cortex were assessed. For each optode, raw light data were extracted and cleaned using median filtering to remove motion artifacts (e.g., head movement) and a low pass filter at 0.1 Hz to remove physiological noise (e.g., heart rate). The 16-channel optode arrangement and experimental set-up are presented in **Figure 1**.

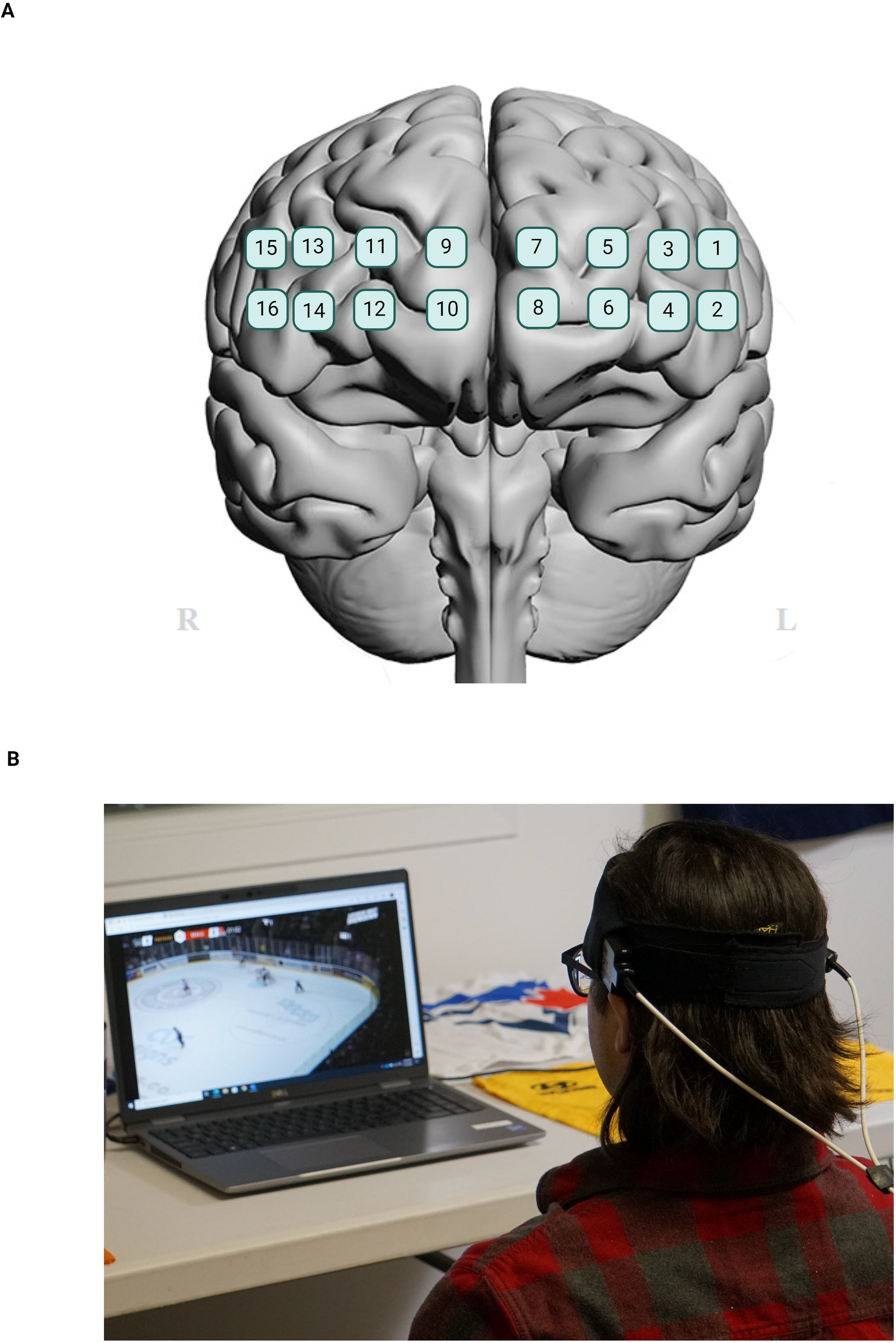

#### Ego Involvement with Sport

To measure EI, an adapted version of a measure described by Kyle et al. (2007) was used. The measure included 15 items regarding the centrality of the target sport (i.e., “[watching ice hockey] is very important to me”), social bonding around the sport (i.e., “[watching ice hockey] occupies a central role in my life”), identity affirmation (i.e., “I identify with the people and image associated with[watching ice hockey]”), and identity expression (i.e., “You can tell a lot about a person by seeing them [watching ice hockey]). Responses to each item were provided using a 5-point scale where 1 = “strongly disagree” and 5 = “strongly agree”. Those scoring +1 *SD* and – 1 *SD* in relation to the overall sample mean were defined as “low involvement” and “high involvement” respectively.

### Signal processing and statistical analyses

Optical data was processed using a median filter followed by a low pass filter. This dual-stage approach has been demonstrated as an effective means of filtering both motion artifacts and physiological noise from fNIRS data (Huang et al., 2022). The filtered optical data was then used to calculate changes in hemodynamic responses using the modified Beer-Lambert law. Channels [3,4,6] were identified as the left lPFC, channels [5,7,8] the left dmPFC, channels [9,10,11] the right dmPFC and channels [12,13,14] as the right lPFC.

During the viewership experience we sampled 12 key moments of interest analyzing the mean change in hemodynamic response in the 10 seconds following a scoring chance or a momentum shift in the offensive end. These key moments consisted of scoring chances and offensive faceoff opportunities (OFO) in which a team gained the puck during a face off in the offensive end. The timing of these key moments is presented in **Table 2**.

**Table 2:** Overview of Key Moments Analyzed During Gameplay.

| Event | Viewing Time (mm:ss) | Period Remaining (mm:ss) |
| --- | --- | --- |
| **Early** |  |  |
| Black Chance 1 | 00:06 | 19:54 |
| Red Chance 1 | 02:22 | 18:22 |
| Red Chance 2 | 04:38 | 16:50 |
| Black Chance 2 | 05:43 | 16:09 |
| **Middle** |  |  |
| Red Chance 3 | 06:22 | 15:30 |
| Black Chance 3 | 09:40 | 15:64 |
| Black OFO | 10:12 | 13:02 |
| Black Chance 4 | 13:08 | 10:10 |
| **Late** |  |  |
| Black Chance 5 | 14:46 | 08:30 |
| Red OFO | 23:44 | 01:02 |
| Red Chance 4 | 24:32 | 00:39 |
| Red Chance 5 | 25:05 | 00:06 |

\*In this table the teams are referred to by their game colors, Red for the Cardiff Devils and Black for the Nottingham Panthers.

The 10 seconds prior to each event was used to establish a local baseline. The 12 key moments used in our analysis are evenly divided into the temporal categories of “Early” (i.e., first four key moments in the period), “Middle” (subsequent four key moments), or “Late” (final four key moments) in order of the viewership experience.

Our statistical analysis was conducted using linear mixed effects models with the lme4 package in R (Bates et al., 2014). Post-hoc analysis was performed using Tukey’s HSD to adjust for multiple pairwise comparisons. We designed our model to examine the relationship between ego involvement and changes in brain activity across brain regions while considering viewing time, and the type of game event as interaction terms. For the fixed effects component of the model, we specified random intercepts for each participant. The final model specification was as follows: HbO ∼ Involvement*Time*Region + Involvement*Type*Region + (1|participant)

## Results

There was a significant two-way interaction between ego-involvement and time *F*(2, 2830) = 5.30, *p* = .005, and a significant two-way interaction between ego-involvement and region *F*(3, 2830) = 3.17, *p* = .024. *Post-hoc* comparisons were used to examine the differences in HbO response between low and high ego-involved individuals across brain regions at different time points. Results (see **Figure 2, panel a**) indicated that within the right dmPFC individuals with high ego-involvement demonstrated significant increases in HbO in response to key moments across the early, middle and late parts of the game *t*(629) = 2.34, *p* = .019, *t*(369) = 2.33, *p* = .020, *t*(369) = 2.52, *p* = .012 respectively. Results also indicated that within the left lPFC in the early moments of the game individuals with low ego-involvement demonstrated a significant decrease in their HbO response *t*(629) = 2.18, *p* = .030.

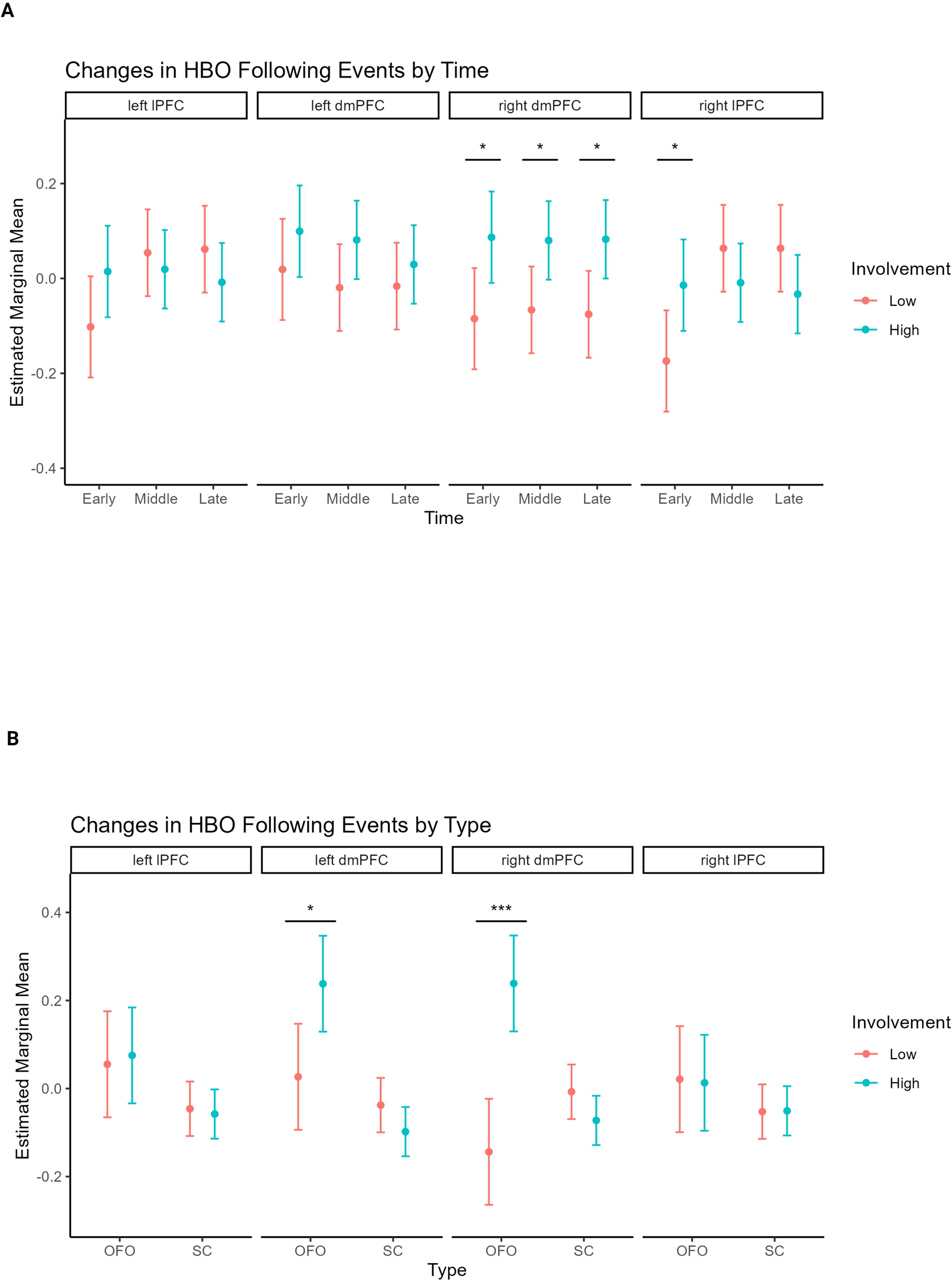

There was a significant three-way interaction between ego-involvement x event type x brain region *F*(3, 2830) = 6.02, *p* < .001. Post-hoc tests were conducted to examine the differences in HbO response across brain regions between low and high involved individuals’ responses to the different types of viewing moments. Results (see **Figure 2, panel b**) indicated that the interaction was driven by significant increases in HbO for individuals with high involvement in both the left *t*(215) = 2.19, *p* = .029 and right *t*(215) = 2.19, *p* = .029 dmPFC during offensive forward opportunities.

## Discussion

The current investigation aimed to examine the feasibility of fNIRS to quantify patterns of brain activation in response to in-vivo sport viewership opportunities, and to evaluate the possibility of moderating effects by ego-involvement. It was hypothesized that fNIRS would prove feasible and provide information about the influence of both critical moments in game play, and ego-involvement, as determinants of cerebral hemodynamics. We observed significant increases in oxygenated hemoglobin (HBO) in the right dmPFC of individuals with high ego-involvement following key moments throughout the course of the game. Comparing the different types of events, it appears that individuals with high involvement were highly responsive to offensive faceoff opportunities as evidenced by increased HBO in both their left and right dmPFC following these moments specifically. Given the important role of the dmPFC in personally relevant social evaluations (Etkin et al., 2011) the increased activation of this structure in individuals with high involvement during sports viewership experiences is theoretically meaningful.

The finding that evaluative processing within the dmPFC differed between high and low ego-involved spectators uniformly across early, middle and late game phases provides validation of the hypothesized evaluative and self-relevance processing function of the dmPFC (Lieberman et al., 2018; Mayer & Lieberman, 2018). That is, it is intuitive that more value-based processing of sensory information delivered via the game play video samples would be evident among participants who were emotionally involved in the sport at the outset of the experiment. Beyond this, when looking at a critical game play moment (i.e., offensive face-offs), there were notably large differences in self-relevance/evaluative processing between high and low-involved spectators.

There are several strengths of the current study. First, few prior studies have examined the sport viewership experience using mobile neuroimaging, and no prior studies have employed fNIRS specifically to our knowledge. From this perspective, the current findings lay the foundation for future applications of fNIRS technology in the sport viewership context and provide a demonstration of feasibility. Second, we took steps to ensure no prior exposure to experimental stimuli was likely by carefully selecting game play sequences from a familiar sport played outside of the host country for the study in question. Finally, we ensured that video stimuli contained a range of prototypical sport event moments, and ensured that the viewer experience was as ecologically valid as possible.

There are also several limitations worthy of mention. First, it is possible that other areas of the cortex (or less superficial levels of the brain in general), including those involved in attentional processes (such as the superior parietal lobule), may be active during specific periods of game play, but these could not be measured in the current study due to optode placement. Second, because the video clips were selected from actual game play, we cannot guarantee that there was no pre-exposure or attitudinal variation that shaped the neural responses differentially for those at different levels of involvement. Finally, the range of events sampled in our visual stimuli was not exhaustive with respect to moments in game play, and there may be other events not captured that more fully reveal differences between those viewers who are involved versus not involved with respect to attention or evaluative processing.

Despite these limitations, the current study is a step forward in the science of sport spectatorship experiences. Objective, continuous (i.e., ongoing throughout the viewership experience), and in situ measures of sport viewership engagement are not typically captured in previous sport consumer investigations (Potwarka et al., 2022). Neuroimaging data may inform design and production elements of more engaging televised sport stimuli for diverse sets of audiences.

## Conclusion

The current study examined the extent to which fNIRS is a viable technology for capturing neurocognitive aspects of the sport spectator experience in an ecologically valid manner, and the extent to which certain key moments in gameplay may be disproportionately influential. We also examined the extent to which pre-existing ego-involvement with the sport viewed might moderate any such effects. Our findings largely supported our hypotheses that fNIRS would be feasible, that some key moments were more important than others, and that ego involvement predicted relatively stronger neural effects, particularly within evaluative and self-relevance processing nodes within the dmPFC. Despite limitations, the observed findings provide an initial confirmation that fNIRS is well-adapted to the sport viewership experience, and that neural signals may reliably differentiate between those who are more versus less highly involved in viewing actual game play sequences, specifically in the context of hockey. Future studies examining the reliability of such effects across different sports would be valuable, as well as experimental manipulation of cortical nodes to unpack temporal precedence and causality.

## References

Ayaz, H., Baker, W. B., Blaney, G., Boas, D. A., Bortfeld, H., Brady, K., … & Zhou, W. (2022). Optical imaging and spectroscopy for the study of the human brain: status report. Neurophotonics, 9(S2), S24001.

Bates, D., Mächler, M., Bolker, B., & Walker, S. (2014). Fitting linear mixed-effects models using lme4. arXiv Preprint 1406.5823.

Etkin, A., Egner, T., & Kalisch, R. (2011). Emotional processing in anterior cingulate and medial prefrontal cortex. Trends in cognitive sciences, 15(2), 85–93.

Ferrari, M., & Quaresima, V. (2012). A brief review on the history of human functional near-infrared spectroscopy (fNIRS) development and fields of application. Neuroimage, 63(2), 921–935.

Funk, D. C., & James, J. (2001). The Psychological Continuum Model: A Conceptual Framework for Understanding an Individual’s Psychological Connection to Sport. Sport Management Review, 4(2), 119–150. 10.1016/S1441-3523(01)70072-1

Furley, P., Bertrams, A., Englert, C., & Delphia, A. (2013). Ego depletion, attentional control, and decision making in sport. Psychology of Sport and Exercise, 14(6), 900–904. 10.1016/j.psychsport.2013.08.006

Grill-Spector, K., Henson, R., & Martin, A. (2006). Repetition and the brain: neural models of stimulus-specific effects. Trends in cognitive sciences, 10(1), 14–23.

Harris, A., Hare, T., & Rangel, A. (2013). Temporally dissociable mechanisms of self-control: early attentional filtering versus late value modulation. Journal of Neuroscience, 33(48), 18917–18931.

Havitz, M. E., & Dimanche, F. (1999). Leisure Involvement Revisited: Drive Properties and Paradoxes. Journal of Leisure Research, 31(2), 122–149. 10.1080/00222216.1999.11949854

Huang, R., Qing, K., Yang, D., & Hong, K. S. (2022). Real-time motion artifact removal using a dual-stage median filter. Biomedical Signal Processing and Control, 72, 103301.

Kam, J. W., Solbakk, A. K., Endestad, T., Meling, T. R., & Knight, R. T. (2018). Lateral prefrontal cortex lesion impairs regulation of internally and externally directed attention. Neuroimage, 175, 91–99.

Kondo, H., Osaka, N., & Osaka, M. (2004). Cooperation of the anterior cingulate cortex and dorsolateral prefrontal cortex for attention shifting. Neuroimage, 23(2), 670–679.

Kyle, G., Absher, J., Norman, W., Hammitt, W., & Jodice, L. (2007). A modified involvement scale. Leisure studies, 26(4), 399–427.

Lieberman, M. D., Straccia, M. A., Meyer, M. L., Du, M., & Tan, K. M. (2019). Social, self,(situational), and affective processes in medial prefrontal cortex (MPFC): Causal, multivariate, and reverse inference evidence. Neuroscience & Biobehavioral Reviews, 99, 311–328.

Madrigal, R. (2006). Measuring the Multidimensional Nature of Sporting Event Performance Consumption. Journal of Leisure Research, 38(3), 267–292. 10.1080/00222216.2006.11950079

Meyer, M. L., & Lieberman, M. D. (2018). Why people are always thinking about themselves: medial prefrontal cortex activity during rest primes self-referential processing. Journal of cognitive neuroscience, 30(5), 714–721.

Schroeter, M. L., Kupka, T., Mildner, T., Uludağ, K., & von Cramon, D. Y. (2006). Investigating the poststimulus undershoot of the BOLD signal—a simultaneous fMRI and fNIRS study. Neuroimage, 30(2), 349–358.

Oppenheim, A. N. (2000). Questionnaire design, interviewing and attitude measurement. Printer Publishers: London, England

Petty, R. E., Cacioppo, J. T., & Schumann, D. (1983). Central and Peripheral Routes to Advertising Effectiveness: The Moderating Role of Involvement. The Journal of Consumer Research, 10(2), 135–146. 10.1086/208954

Petty, R. E., & Cacioppo, J. T. (1990). Involvement and Persuasion: Tradition Versus Integration. Psychological Bulletin, 107(3), 367–374. 10.1037/0033-2909.107.3.367

Pinti, P., Tachtsidis, I., Hamilton, A., Hirsch, J., Aichelburg, C., Gilbert, S., & Burgess, P. W. (2020). The present and future use of functional near-infrared spectroscopy (fNIRS) for cognitive neuroscience. Annals of the New York Academy of Sciences, 1464(1), 5–29.

Potwarka, L., Hall, P. A., Goebert, C., & Ayaz, H. (2022). Immersive Technology and the Virtual Sport Spectator Experience. In M. L. Naraine, T. Hayduk III, and J.P. Doyle (Eds.), The Routledge handbook of digital sport management (pp. 232–244). Routledge. 10.4324/9781003088899-20

Rothschild, M. L. (1984). Perspectives on involvement: current problems and future directions. ACR North American Advances.

OR Rothschild, M. L. (1984). Perspectives on Involvement: Current Problems and Future Directions. Advances in Consumer Research, 11, 216–.

Tompson, S. H., Falk, E. B., Bassett, D. S., & Vettel, J. M. (2019). Using neuroimaging to predict behavior: An overview with a focus on the moderating role of sociocultural context. Social-Behavioral Modeling for Complex Systems, 205–230.

